# Effect of population structure and migration when investigating genetic continuity using ancient DNA

**DOI:** 10.1101/052548

**Authors:** NM Silva, S Kreutzer, C Papageorgopoulou, M Currat

## Abstract

Recent advances in sequencing techniques provide means to access direct genetic snapshots from the past with ancient DNA data (aDNA) from diverse periods of human prehistory. Comparing samples taken in the same region but at different time periods may indicate if there is continuity in the peopling history of that area or if a large genetic input, such as an immigration wave, has occurred. Here we propose a new modeling approach for investigating population continuity using aDNA, including two fundamental elements in human evolution that were absent from previous methods: population structure and migration. The method also considers the extensive temporal and geographic heterogeneity commonly found in aDNA datasets. We compare our spatially-explicit approach to the previous non-spatial method and show that it is more conservative and thus suitable for testing population continuity, especially when small, isolated populations, such as prehistoric ones, are considered. Moreover, our approach also allows investigating partial population continuity and we apply it to a real dataset of ancient mitochondrial DNA. We estimate that 91% of the current genetic pool in central Europe entered the area with immigrant Neolithic farmers, but a genetic contribution of local hunter-gatherers as large as 83% cannot be entirely ruled out.

## Introduction

Neutral genetic diversity in human populations reflects past demographic changes and migrations. While genetic data from contemporary humans has long been the sole source of molecular data used to draw inferences on the evolution and peopling history of their ancestors (e.g., 1), direct evidence from the past has been recently recovered by sequencing ancient DNA (aDNA) from different time periods and geographical regions (2–6). Although full genomes are now published with various coverage (e.g., 7, 8-11), datasets belonging to the same prehistoric “population”, defined either geographically or culturally, have mostly been published for mitochondrial HVS1, such as for the Late Upper Paleolithic and Neolithic era in central Europe and Spain (2, 6, 12). Those data have been used to address questions on the population continuity (PC) through time in the same area, in contrast to a genetic input or replacement due to the arrival of new immigrants (2,13). Ancient mitochondrial population samples have been studied independently for different regions in Europe, and most of them revealed regional genetic discontinuity through time, from prehistoric times until today, meaning that the observed shifts in allele frequencies cannot be explained by genetic drift alone. This conclusion applied to more than 83% of the tests for population continuity applied in Europe (14).

In order to assess if two genetic samples from different time periods but taken in the same geographic area (hereafter called “serial samples”) may be considered as coming from a single population evolving under the sole effect of genetic drift, a model-based test has been developed and applied to mitochondrial data (2). The framework of this test is to simulate serial genetic samples with the same characteristics as the real ones (time period, sample size) issued from one single panmictic population, and to compute an index of genetic differentiation between those samples, usually the fixation index *Fst*. With a reasonable number of simulations (usually several thousand), a distribution of genetic distances between serial samples under the null hypothesis of PC is provided. If the observed genetic distance between the real samples is above 95% of the simulated ones, then the null hypothesis is rejected, meaning that genetic drift alone is not able to generate the differences in genetic diversity between the serial samples. Genetic shift through demographic replacement or migration are the factors commonly proposed to explain such a population discontinuity. For instance, a population discontinuity between Paleolithic hunter-gatherers (PHG) and Neolithic farmers (NFA) from the same area could be interpreted as a large demic replacement of PHG by NFA coming from another area. This test for PC is thus heavily relying on the simulated distribution of *Fst*, and consequently on the underlying model which must be as realistic as possible.

However, while being able to simulate complex scenarios and some very simple levels of population structure, all the programs used so far to generate the null distribution are not spatially explicit and do not consider ancestral migration of genes among neighboring populations (15–17). They thus make the strong assumption that the ancestral lineages of people living today at a given place have always been in the same area in the past. This assumption of panmixia, or near panmixia, through time is questionable given the high mobility of humans. Indeed, studies at micro-regional levels have highlighted the major role played by migrations in partially renewing the local genetic pool during a period as short as a few generations (18–21). Those rare studies incorporating ancient population structure when analyzing aDNA, although in a simplified way, better explain the data (6, 22, 23). Moreover, the incorporation of a spatial component in the analysis of genetic data has already proven to generate insights on human evolution, and even some counter-intuitive results that were undetectable with non-spatial approaches (24–27).

Here, we investigated PC through a spatially-explicit approach by using a modified version of SPLATCHE2 (28), allowing the sampling of genetic lineages both over time and space. We explored the effects of incorporating spatial migration as a fundamental element when testing for PC using aDNA. Moreover, we also investigated how spatial and temporal heterogeneity, which usually characterize aDNA population samples, may affect the analysis. Indeed, one big difference that characterizes ancient and modern datasets is the temporal and geographic heterogeneity present in the population samples. Because of the scarcity of aDNA, lineages quite distant in time and space are often grouped together based on cultural or geographic criteria. For instance, the hunter-gatherers sample published by (2) encompasses ancient lineages with a range of dates from 2,250 calBC to 13,400 calBC and a geographic range including samples as distant as 3,000 km. By contrast, modern population samples are much more homogeneous, in terms of time but also of geographical scale. If temporal heterogeneity within population samples may be assumed in the programs used so far to investigate PC, such as BSSC (15) and FastSimCoal (16), spatial heterogeneity cannot. Our spatially-explicit approach thus constitute a solution to cope with these two dimensions simultaneously. In addition, our approach allows to test for partial PC, contrary to previous approaches which only tested for full (100%) PC. We applied our spatially-explicit approach to a real dataset of PHG and NFA samples from central Europe, in order to estimate their relative genetic contribution to the modern gene pool of this area.

## Materials and Methods

### Spatially-explicit simulation of ancient DNA

A modified version of the program SPLATCHE2 (28) allowing the sampling of lineages at different time points was used, making it possible to reconstruct the coalescent tree for genetic samples of different ages. This version improves model-based approaches previously used to test for PC with ancient DNA (15, 16) by considering the spatial dynamics of genes in a spatially-explicit context. Moreover, it allows testing for partial population continuity using the two population layer mode of SPLATCHE and the admixture rate parameter γ (29).

The framework comprises as a first step the simulation of a population expansion in a grid of demes exchanging migrants in a stepping-stone fashion. The area could be previously either empty or already occupied by a local population. In a second step, a coalescent reconstruction is performed in order to generate mitochondrial genetic diversity in samples of different ages and locations drawn from the simulated population. The genealogy of simulated lineages is reconstructed conditional to the density and migration rate calculated during the first step (30). The genetic diversity of those lineages is simulated by distributing mutations on the coalescent tree, using a mutation rate μ for a DNA sequence of a given length, which represents, in this study, the mtDNA HVS1 region. See the original description of SPLATCHE for more details on the algorithms (28).

We used two spatially-explicit frameworks:

(1) a virtual square map in which we explored the influence of the spatial component when investigating the relationships between serial samples;
(2) a realistic European map, which was used to estimate the relative genetic contribution of local PHG and incoming NFA to the genetic pool of Central European populations.

## Simulations on a square map

### Comparison between spatial and non-spatial models when investigating PC

The assumption of genetic continuity between two serial samples implies that they descend from the same population and that the genetic differences between them are only due to genetic drift and sampling (non-spatial model, NSP) as well as gene flow with neighboring populations (spatial models, SP). Where there is continuity, the population evolves simply by continuously exchanging a small amount of genes with its neighbors but without a large genetic input or replacement from another differentiated gene pool. To represent PC, we thus simulated the expansion of one population in a grid of demes exchanging migrants (SP); or a simple demographic growth in a single deme (NSP). For SP, we simulated the growth of a population of 100 individuals during 2,000 generations (~50,000 years for humans) from the center of a square map of 2,500 demes (50 × 50), as described in (27), with a carrying capacity *K* of 500 individuals each. A logistic equation controls the population growth at the deme level with a growth rate *r*. This spatial model, SP, was compared to a non-spatial version, NSP, made up of only one deme with parameters adapted for the purpose of comparison with the spatial model (see Figure 1): no migration (*m* = 0), the population growth was equal to 0.012 in order to reach *K* at the end of the simulation, with *K* equal to the product of the number of demes and *K* in the spatial model (500 individuals × 2,500 demes). Note that all parameters are constant for the whole duration of the simulation if not indicated otherwise.

**Figure 1.**
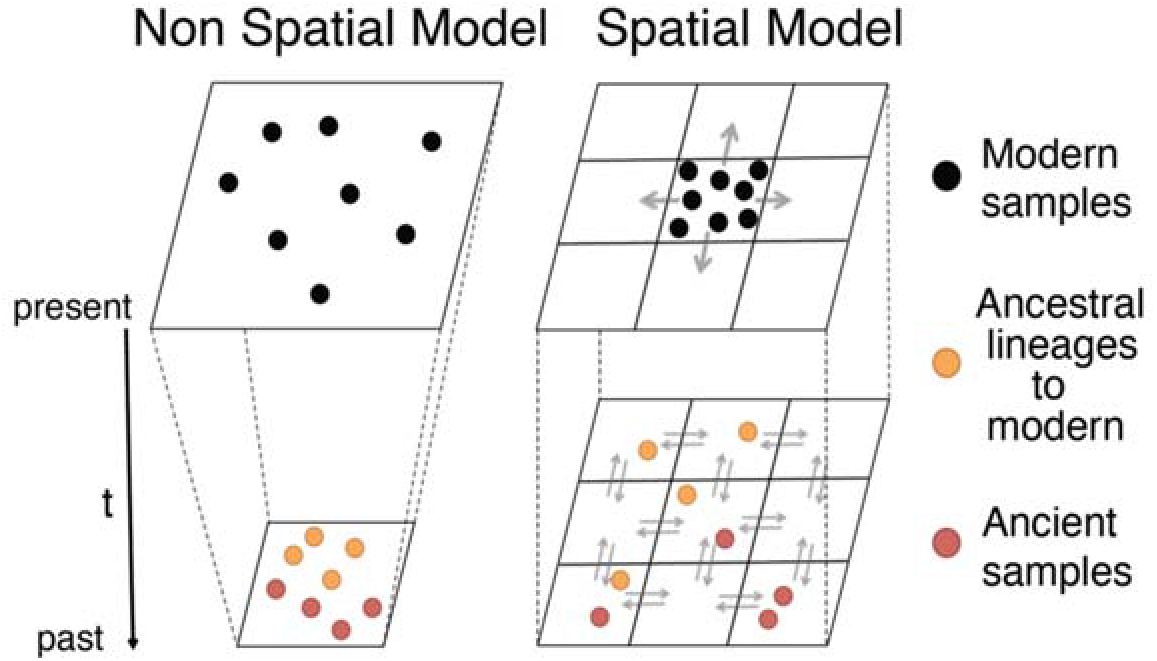
Schematic representation of lineages simulated under non-spatial (NSP) and spatial models (SP) of population continuity. The NSP representation stresses the geographical constraint imposed in such models, with the individuals staying in the same geographical location during the whole simulation. This constraint is absent in a spatial model, with migration and population structure adding an important realistic element.

For the spatial model, we performed several series of simulations with various migration rates *m* (values 0.005, 0.01, 0.05 and 0.1) in order to explore the effect of population structure on the genetic differentiation between two serial samples drawn from a continuous population. *Nm* is the migration rate *m* multiplied by the size *N* of the deme, and defines the number of migrants *Nm* that are distributed in the demes located adjacently (at maximum four: North, South, East and West) at each generation. A high *Nm* means that the population is large and interconnected while a small *Nm* means that the population is small and isolated. For all scenarios, *Nm* is kept constant throughout the simulation, but we also simulated a Neolithic scenario where *Nm* was changed from 5 to 50, 400 generations before present to reflect changes following the Neolithic transition (31). This was done modifying *m* in all demes at the same time. We also investigated the effect of various combinations of *K* (100,150 and 200) and *m* resulting in the same *Nm* of 5 or 50.

For each simulation, two groups of 30 mtDNA lineages, called “population sample” hereafter, were sampled in the center of the map at two different times: a modern population sample at present (after 2,000 generations simulated) and an ancient population sample 400 generations before the present (after 1,600 generations simulated), which corresponds approximately to the Neolithic time (~10,000 BP). A mutation rate μ = 3.3×10^−6^ for DNA sequence of 300 bp was simulated in order to approximate mtDNA diversity in European populations (27).

A measure of genetic differentiation, *Fst*, between those samples was computed using the software Arlequin 3.5 (32), providing a null distribution of genetic differentiation under the hypothesis of PC. In order to better explore the stochastic nature of the coalescent processes, 10,000 independent simulations were performed for each scenario (combination of parameters). The final results are presented as a distribution of *Fst* for each scenario. We used the same software to compute the gene diversity within each sample (modern *H*_*mod*_ or ancient *H*_*anc*_). We also computed the average coalescent time 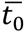 between lineages within population samples (ancient 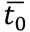*anc* and within modern 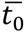*mod* and the average between them) and the average coalescent time 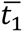 between lineages belonging to different population samples (between modern and ancient).

### Simulation of spatial and temporal heterogeneity within ancient population sample

In two other series of simulations in the square world, we explored the effect on the comparison between serial samples of spatial and temporal heterogeneity between aDNA sequences belonging to the same ancient population sample. We chose two different levels of population structure reflected by *Nm* values of 5 and 50, and varied the heterogeneity within the ancient dataset, dividing the 30 ancient lineages into five groups as shown by Figures 2A and 2C and described below. Although the ancient dataset was divided into five groups of lineages, it was considered as a single ancient population sample when computing *Fst* with the modern sample, to reflect the kind of grouping usually made with real data(2,5)

**Figure 2.**
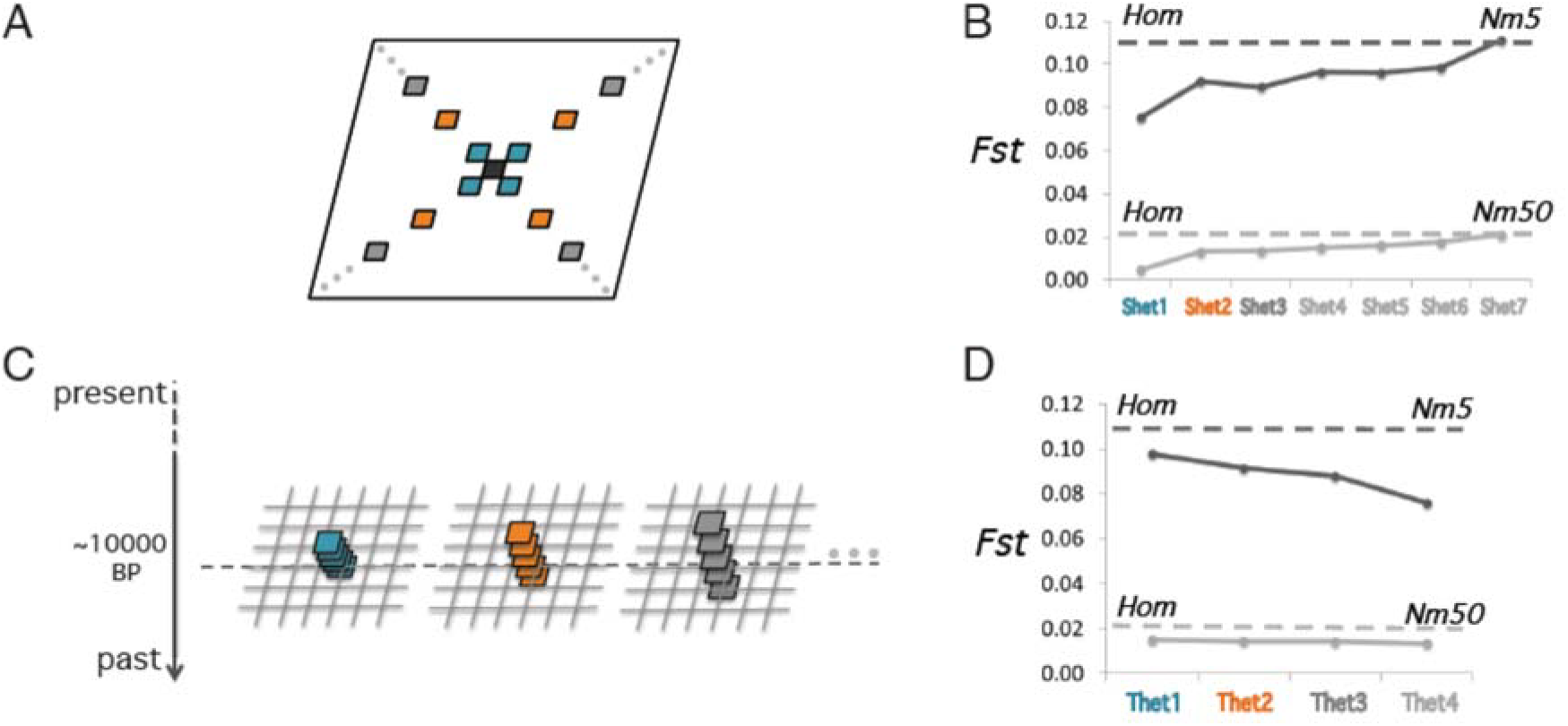
Effect of spatial and temporal heterogeneity within ancient population sample when computing genetic distances between ancient and modern samples. Modern lineages are always drawn in one single central deme. A), ancient lineages are drawn by groups of 5 in 5 different demes, separeted geographically with increasing distance from the center, along the two diagonals. Shet7 is the most dispersed and Shet1 the least. B) The *Fst* is calculated between the group of modern lineages and the group of ancient lineages considered as one population sample. The *Fst* values are displayed for all sampling schemes, in comparison with a homogeneous one (Hom, dotted horizontal line) for *Nm* equal to 5 (dark grey) and 50 (light grey). C-D) The same principle is applied for the variation in temporal sampling with Thet4as the most variable in terms ofagesandtheThet1 the least.

For spatial heterogeneity, we tested seven different spatial configurations, all with one group of five lineages in the deme located in the center of the map and four other groups of five lineages each taken in four demes along the diagonals of the square area – with increasing distance from the center deme (sampling “Shet1” to “Shet7”).

For temporal heterogeneity, we tested four temporal configurations, with all lineages taken in the central deme but at different times. One group of five lineages is always drawn 400 generations before the present, but two other groups of five are taken more recently, and the 2 others at more ancient points. The temporal distance between groups increases from “Thet1” to “Thet 4” scenarios.

### Simulations on a European map

We then implemented our approach on a digital map of Europe divided in demes of 100×100 *km* in order to investigate the genetic influence of the Neolithic transition in Central Europe using the two-layers model designed by Currat and Excoffier (29). The first layer represents the expansion of a PHG population of 100 individuals starting 1,600 generations ago (~40,000 years) from southern Middle East. Each PHG deme has a carrying capacity (*K*) of 100 individuals corresponding to 0.06 individuals/km^2^ (33, 34). The migration rate (*m*) and growth rate (*r*) were calibrated to 0.15 and 0.2, respectively, using 500 generations as the time of colonization of Europe by *Homo sapiens*, following Currat and Excoffier (29). The second layer represents the Neolithic expansion starting from the Fertile Crescent 400 generations before the present (~10,000 years). The source NFA population is made up of 100 individuals. Each NFA deme has a carrying capacity *(K)* of 1,000 individuals which corresponds to the maximum density estimated for the LBK *(LinearBandKeramic)*, equal to ~0.6 individuals/km^2^ (31). All density and carrying capacity values are given in haploid effective size. For the Neolithic layer, the migration rate *(m)* and growth rate (*r*) were calibrated to fit the dates of the Neolithic samples under study. A migration rate *m* = 0.4 and a growth rate *r* = 0.53 were estimated, corresponding to a speed for the spread of farmers in Europe equal to 1.13 km/y, in accordance with what was estimated by (35). The gene flow between the two layers depends on the parameter γ which represents the proportion of contact between individuals from the two layers resulting in admixture). A γ equal to its minimum 0.0 means no admixture between HG and NFA and its maximum 1.0 means full admixture. It also represents the assimilation rate or the proportion of hunter-gatherers adopting farming after contact with farmers.

In order to cope with the long persistence of hunter-gatherers in central and northern Europe (13), we modified the original model by lengthening the cohabitation period in central Europe for 200 generations, until 5,000 years ago. This modification allows us to sample in the hunter-gatherer layer at the time corresponding to the youngest hunter-gatherer sample of our dataset (table 1), otherwise the deme would be empty, all PHG having already been replaced by NFA.

**Table 1.** Characteristics of the ancient mitochondrial samples used in the analyses, including temporal and geographic information.

| *Group* | *Abbreviation* | *Sample Count* | *Site* | *Geographic Region* | *Sample Age (cal BCE)* | *Archaeological Context* | *Latitude* | *Longitude* | *Individuals per site* | *references* |
| --- | --- | --- | --- | --- | --- | --- | --- | --- | --- | --- |
| **Hunter Gatherer Central Europe** | **PHG** | 19 | Hohler Fels | Germany | 13400 | paleolithic | 48.38 | 9.76 | 1 | (2) |
|  |  |  | Bad Dürrenberg | Germany | 6990-6706 | mesolithic | 51.30 | 12.07 | 1 | (2) |
|  |  |  | Hohlenstein-Stadel | Germany | 6743 | mesolithic | 48.55 | 10.17 | 2 | (2) |
|  |  |  | Oberkassel | Germany | 11641-11373 | paleolithic | 50.71 | 7.17 | 1 | (49) |
|  |  |  | Blätterhöhle | Germany | 9210 - 8638 | mesolithic | 51.36 | 7.55 | 5 | (13) |
|  |  |  | Loschbour | Luxembourg | 6147-6047 | mesolithic | 49.77 | 6.24 | 1 | (10) |
|  |  |  | Blätterhöhle | Germany | 3922 - 3359 | neo FHG | 51.36 | 7.55 | 8 | (13) |
| **Early Neolithic Central Europe (LBK)** | **NFA** | 99 | Derenburg | Germany | 5500-4775 | LBK | 51.87 | 10.91 | 20 | (4,5,48) |
|  |  |  | Halberstadt-Sonntagsfeld | Germany | 5500-4775 | LBK | 51.90 | 11.06 | 31 | (3,5) |
|  |  |  | Karsdorf | Germany | 5500-4775 | LBK | 51.28 | 11.65 | 23 | (3,48) |
|  |  |  | Naumburg | Germany | 5500-4775 | LBK | 51.15 | 11.81 | 4 | (3) |
|  |  |  | Oberwiederstedt 1, Unterwiederstedt | Germany | 5500-4775 | LBK | 51.67 | 11.53 | 8 | (3,5) |
|  |  |  | Eilsleben | Germany | 5000 | LBK | 52.15 | 11.22 | 1 | (5) |
|  |  |  | Schwetzingen | Germany | 5500-4775 | LBK | 49.38 | 8.57 | 4 | (5) |
|  |  |  | Vaihingen | Germany | 5500-4775 | LBK | 48.93 | 8.96 | 1 | (5) |
|  |  |  | Seehausen | Germany | 5500-4775 | LBK | 51.33 | 11.13 | 1 | (5) |
|  |  |  | Flomborn | Germany | 5500-4775 | LBK | 49.69 | 8.15 | 6 | (5) |

In order to estimate the admixture rate γ between PHG and NFA during the Neolithic transition in Central Europe, we applied an Aproximate Bayesian Computation (ABC) approach using the ABCtoolbox (36). To cope with the uncertainty of the six demographic parameters (migration rate, growth rate and carrying capacity for both layers), we defined prior distribution centered on the values estimated from the literature and described above. The parameter γ was sampled from a lognormal prior distribution because variation of low γ values is more influential than variation of high values regarding the amount of PHG contribution to the NFA layer and we thus want to explore in more details lower γ values. Hundred and sixty thousand simulations with distinct combinations of parameters drawn from the prior distributions (Table 2) were done and used to calculate the *FST* between the two samples (PHG and NFA) in central Europe. The 1,000 combinations of parameters that produced *Fst* values the closest to the observed *Fst* were retained and used to estimate the most plausible parameter values. We mainly focused on the estimation of our parameter of interest: the admixture rate γ, while the other parameters served to test the robustness of the estimation of γ.

**Table 2.**
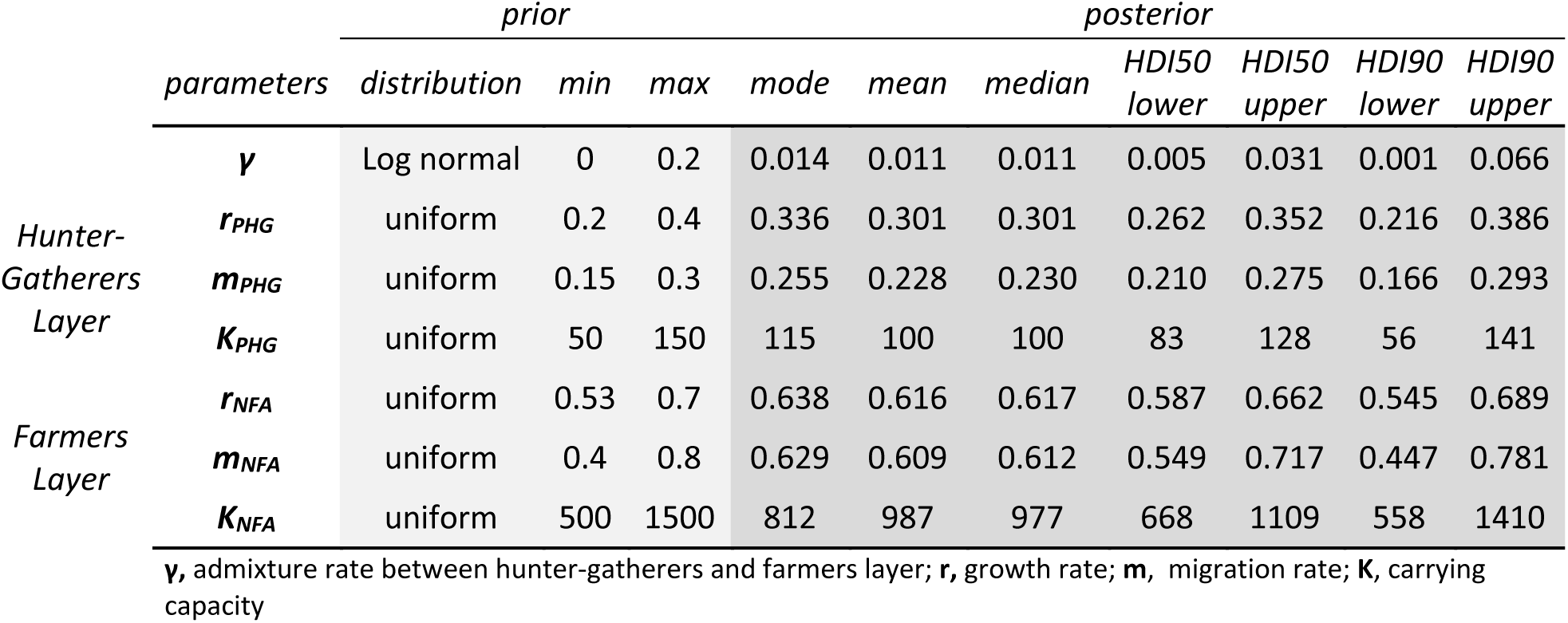
Characteristics of prior and posterior distributions for the model’s parameters used in the ABC estimation procedure.

From the estimated γ values, we were able to estimate the proportion of genetic replacement in Central Europe due to the incoming farmers following the same procedure than Currat and Excoffier (29). We ran the model with the combination of the best parameters and sampled 25 locations in central Europe where the proportion of genes coming from the source population of NFA was calculated. We did it with two values of γ: the point estimate and the value corresponding to the upper limit of the 90% highest density interval (HDI).

We reproduced by simulation mitochondrial samples identical to real data in terms of lineage number, location and age (Table 1 and Figure 3). We reproduced the real dataset by simulating 99 Farmers and 19 Hunter-gatherers mitochondrial HV1 DNA sequences of length 344 (5 additional positions were left out due to too many missing data in the whole dataset), using a mutation rate of 0.0000075 mutations/generation/site.

**Figure 3.**
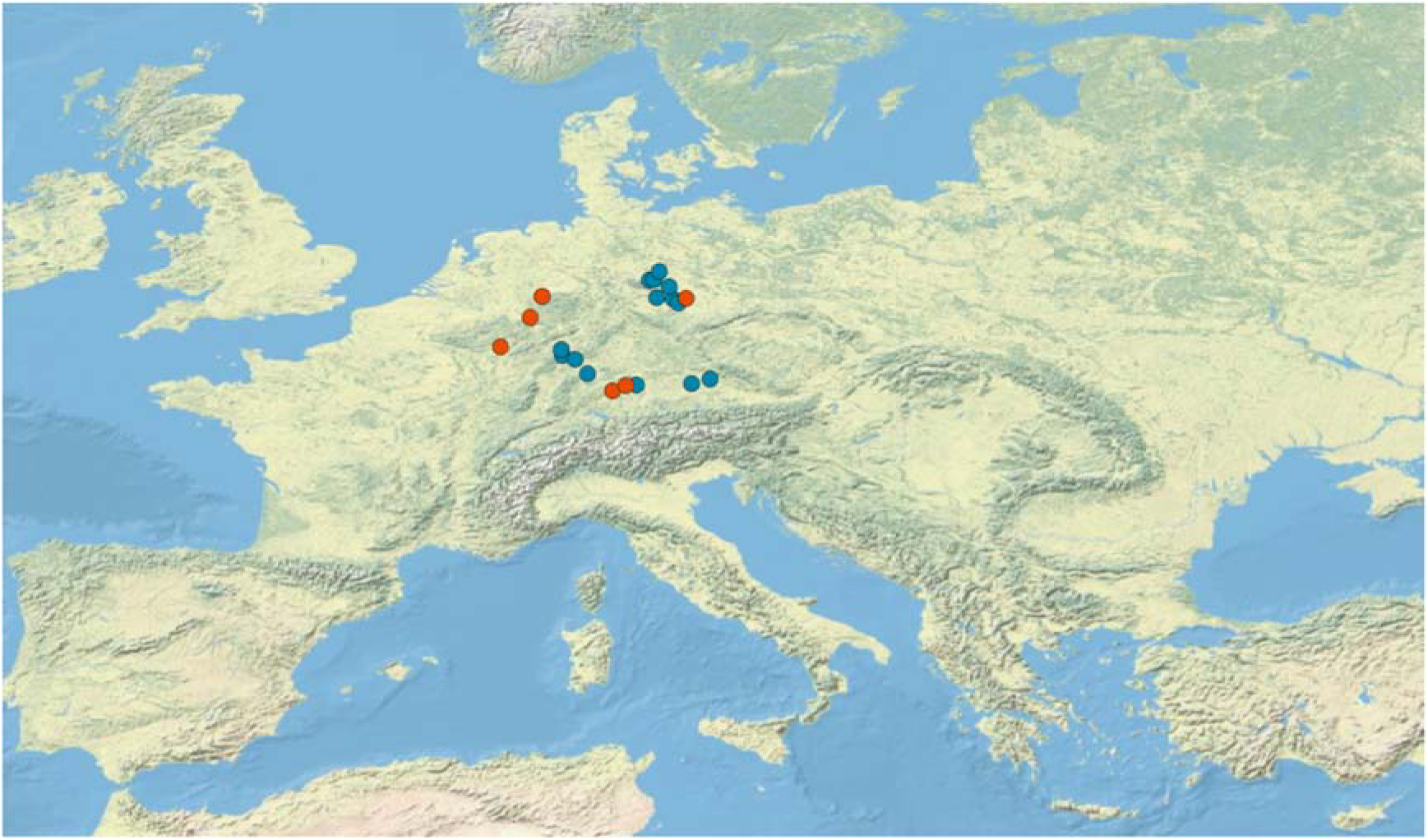
Geographical locations of the Central Europe Hunter gatherers (orange) and Farmers (blue) mtDNA sequences used in this study. The dashed-line rectangle represents the longer cohabitation zone between PHG and FA imposed in the model (map source: US National Park Service).

## Results

### Comparison between spatial and non-spatial models when investigating PC

In comparison to a non-spatial model, all spatial scenarios give larger *Fst* mean and variance between the two serial samples simulated whatever the value of *Nm* (Figure 4A and Table 3). Among the spatial scenarios, the *Fst* and its variance tend to increase with the level of population structure. In other words, the *Fst* increases when *Nm* decreases. For the “Neolithic” scenario, with a shift from *Nm* 5 to *Nm* 50, 400 generations before the present (~10,000 years), intermediate values of *Fst* between *Nm* 5 and *Nm* 50 are found but slightly closer to *Nm* 5, indicating that the “ancient” *Nm* is more influential than the recent one and showing that demographic dynamics through time affect the simulated genetic diversity. This is also visible with gene diversity which is more similar to the ancient gene diversity than to the modern one (Table 3). Regarding the effect of varying *K* and *m* for a same *Nm* value (5 and 50), the *Fst* between the two samples slightly increases with the decrease of *K* (Figure S1). The genetic diversity in both, ancient and modern samples, slightly decreases with *K* and the effect is stronger for *Nm 5*.

**Figure 4.**
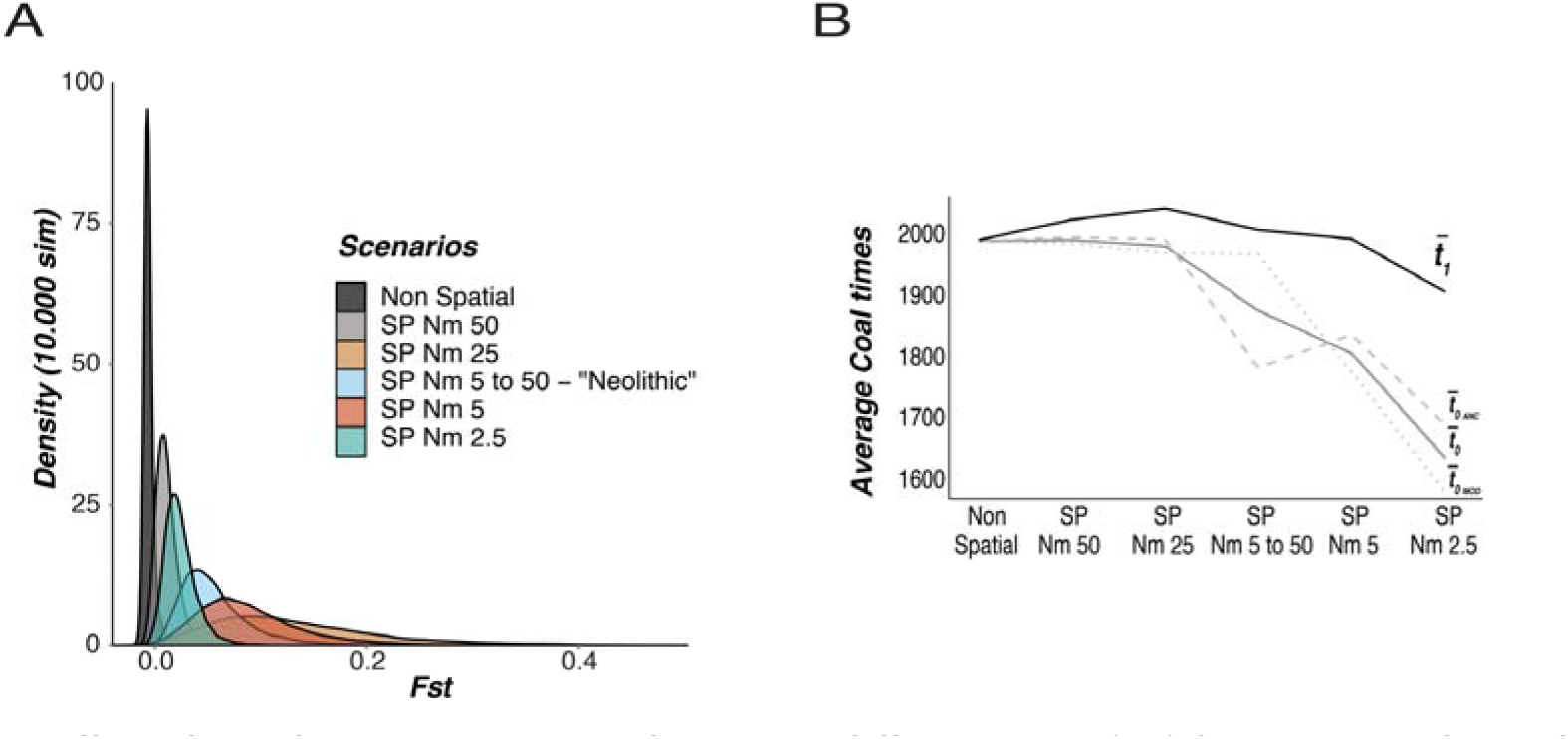
Effect of population structure on the genetic differentiation *(Fst)* between serial samples. From a non-spatial scenario (1 single deme) to spatial scenarios with increasing levels of population structure between 2,500 interconnected demes (SP, with structure increasing proportionally with decreasing *Nm*). A) *Fst* distribution for the scenarios tested and B) evolution of average coalescent times at the intra population 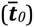 and inter-population levels 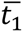.

**Table 3.**
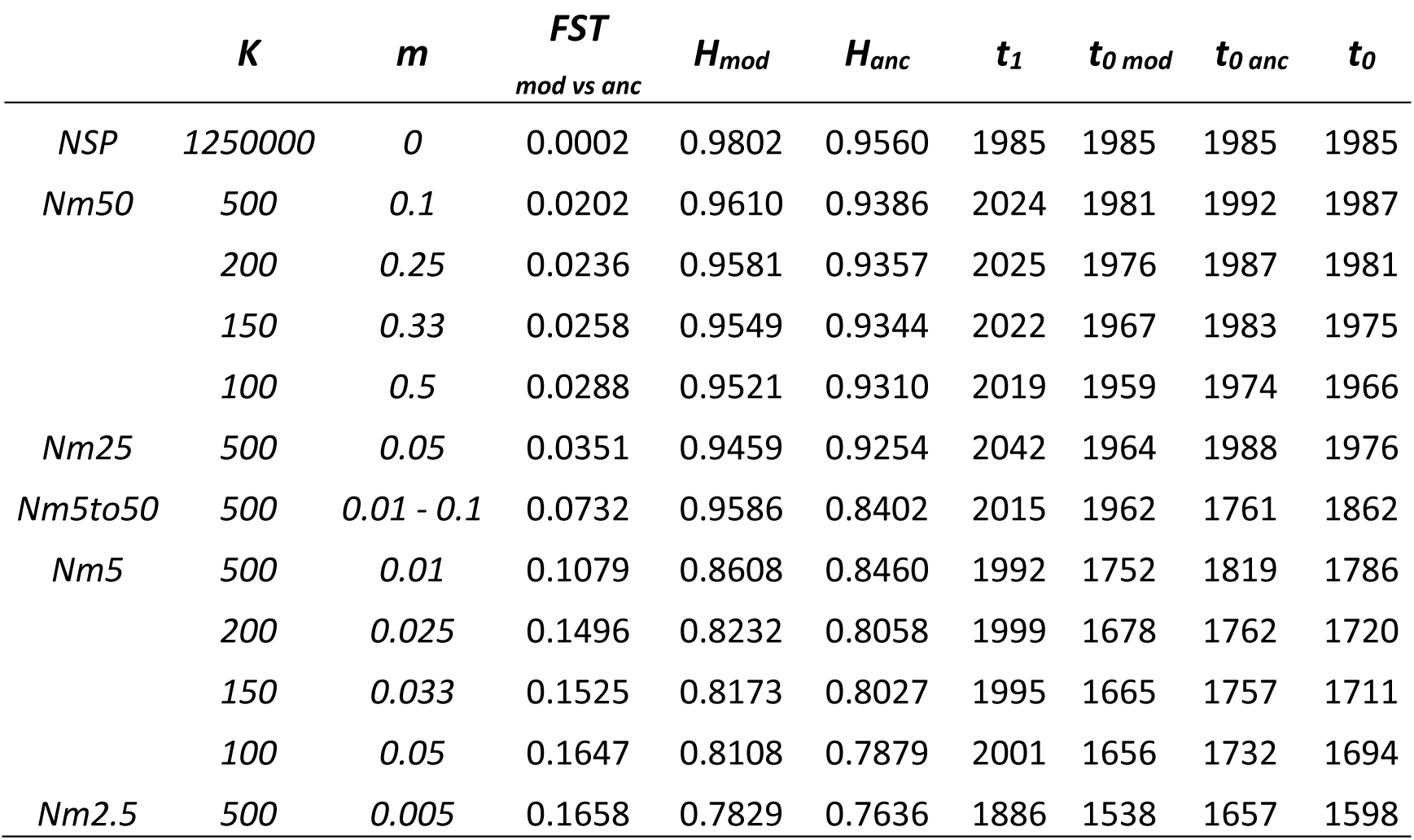
Statistics calculated for the different scenarios used to investigate the effect of population structure and migration when testing for genetic differentiation between serial samples.

For the same set of simulations, we computed the average coalescent time between lineages within population samples 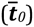 and the average coalescent time between lineages belonging to different population samples 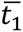 (Figure 4B and Table 3). In a non-spatial scenario, 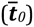 and 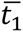 are similar because the coalescent tree is “star-like” and most of the coalescence occurs close to the root, at the onset of demographic expansion when the population size is small with modern and ancient lineages sharing common ancestors. For spatial scenarios, decreasing *m* favors earlier coalescent events between lineages from the same population sample, as shown by diminishing 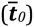 (Figure 4B and Table 3). The average coalescent time 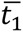 between samples also diminishes but at a smaller rate. Increasing the population structure thus results in a higher genetic homogeneity within samples, and consequently more genetic differentiation between samples, as measured by the *Fst*, which is proportional to 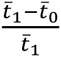 (37). 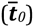*anc* is slightly more affected than 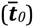*mod*. The particular cases of identical *Nm* with varying *K* show that the decrease in 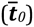 is more pronounced than the decrease in 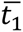, especially when *Nm* is small.

### Influence of temporal and spatial heterogeneity within the ancient sample

We then varied the spatial or temporal heterogeneity within the ancient population sample, both in a small-structured population (*Nm*5, i.e. Paleolithic hunter-gatherers) and in a larger population with more gene flow (*Nm*50, i.e. Late Neolithic famers or historical populations), and computed the *Fst* between the ancient and the modern sample, taken from a single deme. Our results show that spatial and temporal heterogeneity within the ancient sample decrease the *Fst* compared to a homogeneous sampling. In addition, *Fst* tends to increase with spatial heterogeneity (Figure 2B) and to decrease with temporal heterogeneity (Figure 2D). Overall, these results show that variation in time and geographic locations of lineages within the population samples influences the inter-population genetic relationships, and this is stronger when *Nm* is smaller.

### Genetic effect of the Neolithic transition in Central Europe

We estimated an admixture rate γ between PHG and NFA in central Europe of 0.01 with a high density interval (HDI) varying from 0.001 to 0.066 (Figure 5 and Table 2). This result means that around 1% of the contacts between PHG and NFA resulted in the adoption of farming by PHG or to the birth of a child in the farming community. All the best parameter values are given in table 2. Our model reproduces robustly the observed *Fst* as revealed by the marginal density *p*- value of 0.991 and the posterior predictive check (Figure S2). The quantile and HDI distributions are quite uniform as expected for an unbiased estimation of γ (Figure S3). When we translated the values of γ into the genetic input of immigrant NFA, we get 91% with an HDI interval between 17% and 100%. It means that the most likely local PHG genetic contribution is around 9% of the current modern genetic pool but a much higher contribution (up to 83%) cannot be ruled out.

**Figure 5.**
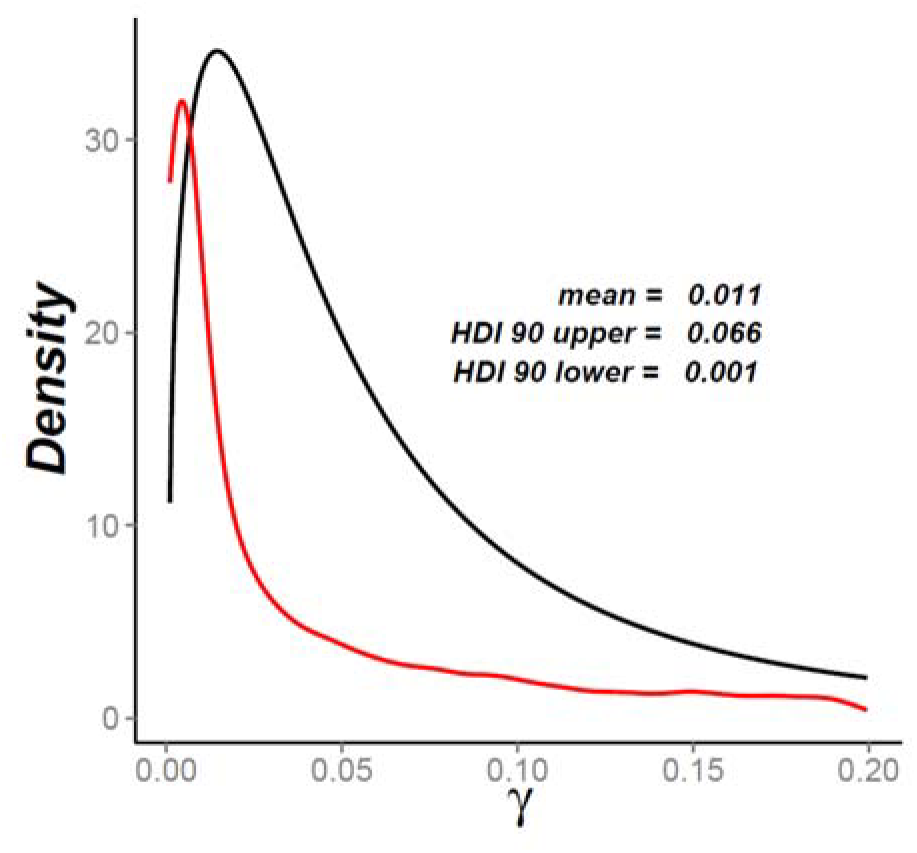
Representation of the prior (in red) and posterior (in black) distributions for the estimated parameter γ from *Fst*.

## Discussion

### Including spatial dynamics of genes when investigating population continuity

To test if two serial genetic samples drawn from the same area at different time are derived from a continuous population, a summary statistics measuring the genetic differentiation between them is computed. Then, this real value is compared to the distribution of equivalent statistics computed on virtual genetic data modeled under a null hypothesis of PC. However, the PC test used to date was non-spatial in the sense that it simulated a single deme without considering any population structure or migration, which are known to be important factors in human evolution (27, 38–40). We demonstrate here that these two elements affect significantly the test for PC as they increase the *Fst* obtained under the null hypothesis to which the real data are compared. The comparison between simulated and real data is made by computing a *p*-value, which is the proportion of simulated *Fst* values bigger than the empirical value, following the procedure of (2). The model of PC is rejected if the *p*- value is below a 5% threshold. So a larger observed *Fst* is needed to reject the null hypothesis of PC when the spatial dynamics of genes are considered.

When using a spatially-explicit modeling framework, the amount of genetic differentiation between samples, *Fst* increases inversely to the composite parameter *Nm*, which represents the amount of gene flow between populations (Figure 4A). The increase of *Fst* together with the reduction of *Nm* is due to a larger number of coalescent events between lineages belonging to the same population sample (either ancient or modern, *t*_*0*_ in Figure 4B), while the average number of coalescent events between lineages belonging to different samples are much less affected (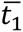in Figure 4B). Two effects occurring during the “scattering phase” (41, 42) are involved in the decreases of 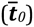. First, lower *m* decreases the probability of emigrating; lineages tend to stay longer in the deme where they have been initially sampled, and are more likely to undergo a coalescent event. Second, lower *N* increases the probability of a coalescent event between two lineages located in the same deme, as it is proportional to *1/2N*. If lineages belonging to the same sample have more coalescence together (lower 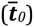) they will tend to be more similar to each other, and more different from one another, on average than the lineages of other samples. The effect of *Nm* on 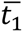 is much less, because going backward in time, as soon as lineages have left the initial deme, corresponding to the “collecting phase” (41, 42), they have a very low probability of encountering another lineage in the grid of demes, whatever the value of *Nm*. Most of the remaining coalescent events occur close to the onset of the expansion when the spatial distribution of the population and its size are low. Testing for PC is more conservative and thus suitable using the spatially-explicit approach than the non-spatial one, because a larger observed *Fst* is necessary to reject the null hypothesis.

### Heterogeneity within ancient samples

Taking into account the spatial and temporal heterogeneity within ancient population samples also affects the *Fst* simulated between ancient and modern samples. When ancient lineages are taken from different demes or at different time periods, *Fst* decreases between serial samples in all cases, while being always below the NSP value (Figures 2B and 2D). This can be explained by the fact that the ancient lineages are initially spread in various demes and have thus a smaller probability of having a coalescent event between them during the scattering phase, thus increasing *t*_*0*_. Consequently, the proportion of coalescent events occurring close to the root of the tree increases with sampling variance and the differentiation between samples diminishes. It is worth noting that the increase of spatial and temporal heterogeneity have different effects. While *Fst* tends to decrease with temporal heterogeneity, it tends to increase with spatial heterogeneity, but in a model that includes spatial or temporal heterogeneity, it still stays below the *Fst* measured using a homogeneous population sample. This result shows that not only the variation in time but also the geographic variation matters when testing for serial inter-population genetic relationships. *Fst* rises with spatial heterogeneity (but still below homogeneity) within the ancient sample because the location of the ancient lineages covers a larger area, on average, than the location of the modern lineages (all from the central deme). Consequently, because they are distant from the center, ancient lineages have a higher probability of having coalescence among them than with modern lineages. Temporal heterogeneity has the opposite effect. The geographical origin of all ancient lineages is the location of the modern lineages (central deme) but when heterogeneity increases, some ancient lineages get closer in time to the modern ones, although the mean age of both population samples stays the same. This closeness in time of some ancient and modern lineages increases their probability of coalescence and consequently diminishes the *Fst* between them.

### Spatiality especially matters when analyzing old prehistoric samples

The effects of population structure and sample heterogeneity are thus additive with some level of variation depending on the amount of heterogeneity. Importantly, the effects of both are stronger in populations with smaller gene flow among sub-populations or deme size (small *Nm*) than in populations with large *Nm* (Figures 4A and 2B and D). This is important because the further back in time we go, the smaller and more isolated the prehistoric populations were (e.g., 43). For instance, *Nm* estimated in current hunter-gatherer populations is < 10 (44). This suggests that taking into account the spatial distribution of lineages is particularly important when investigating PC using prehistoric populations (e.g., Paleolithic). In order to evaluate the effect of population size varying with time, we simulated a “Neolithic” scenario where *Nm* is changing from a low value of 5 to a larger value of 50 at a time corresponding roughly to the Neolithic transition, approximately 10,000 years ago. We used a prehistoric population sample at the first generation after this change (the beginning of the Neolithic era). The results show that despite a high *Nm* (= 50) both at the time of the prehistoric sample, taken at the beginning of the Neolithic era, and the modern one, the distribution of *Fst* measured is between the distributions obtained for *Nm*=5 and *Nm*=50, but closer to the former. Thus, even if current gene flow and/or density is high, *Fst* is strongly affected by the ancient combination between gene flow and density *(Nm)* just before the time of ancient sampling (in this scenario, one generation before sampling) because it affects diversity. The test for PC is thus strongly affected by ancient *Nm*, which means that taking into account spatial structure is especially important in most situations when old samples are analyzed.

### Genetic effect of the Neolithic transition in Central Europe

Except in specific situations such as isolated islands, a population never evolves for centuries without any genetic input from outside. In most cases, there is at least some gene flow from neighboring populations, and this can extend to a full genetic replacement in extreme cases of extermination caused by warfare or disease. The question of population continuity or discontinuity does not have a simple, binary answer. In reality, population continuity may be partially disrupted by migration and result in various amounts of genetic input (or replacement). We thus took advantage of our approach to estimate the amount of genetic replacement during the Neolithic period compatible with ancient mtDNA data in central Europe, by simulating two layers of population in SPLATCHE2, one representing PHG and the other one representing NFA, with various levels of admixture between them (parameter γ). We estimated a genetic contribution of Neolithic farmer immigrants, possibly from south-eastern Europe (17, 45, 46), to the modern Central European genetic pool of 91%, the rest being transmitted by local PHG adopting farming. Our result may explain the genetic shift reported between the Late Upper Paleolithic and the Neolithic in central Europe (2) and the relative contribution of two divergent ancestry components detected by genomic studies (10): the early European farmer (EEF) and the west European hunter-gatherer (WHG). Note that we did not consider possible Yamnaya migration from the Pontic steppes in our model (47). While full population continuity (100% genetic contribution of PHG to the Neolithic community) can be excluded, thus confirming the previous estimation by (2), a PHG genetic legacy as high as 83% cannot. Our estimations give orders of magnitude but show that the method is valuable when investigating past genetic history of human populations. The estimation is done for mitochondrial DNA and thus is valid for the female line only. The same kind of estimation applied to Y chromosome data (if applicable in the future), may give a different answer (23). Moreover, our future aim is to extend this approach to genome wide data in order to precise the estimation.

## Conclusion

Overall, our results underline the need to consider the spatial dynamics of genes when analyzing ancient population samples, because the effects of the various temporal and spatial processes in action (geography, migration, sampling) are complex. Indeed, considering gene flow and population structure in the model increases the expected genetic difference between serial samples compared to previous non-spatial approaches, while spatial diversity and temporal diversity within population samples also affect this differentiation in diverse ways. We also demonstrated that by using a spatially-explicit model when testing for population continuity is valuable when analyzing samples drawn from small and isolated populations. A spatially-explicit model is thus especially suited to the study of prehistoric populations. Despite we simulated mitochondrial diversity, our main results are valid for any kind of comparison between serial molecular samples and thus may be used to help interpreting results obtained from genomic data. Applied to a real dataset from central Europe, our approach confirms a partial genetic shift between Late Upper Palaeolithic and the Neolithic (around 91%), but cannot totally reject a local hunter-gatherer contribution as high as 83%.

## Authors’ contributions

NMS carried out the simulations and analyses. MC conceived, designed and coordinated the study. NMS and MC interpreted the results and drafted the manuscript. SK and CP compiled the data and contributed to the writing of the manuscript. All authors gave final approval for publication.

## Acknowledgements

Computations were performed in the High Performance Computing (HPC) cluster baobab.unige.ch. We thank Laurent Excoffier, Joachim Burger and Daniel Wegmann for stimulating discussion on the subject.

## References

1. Cavalli-Sforza LL, Menozzi P, Piazza A. The history and geography of human genes. Princeton, N.J.: Princeton University Press; 1994. xi, 541, 18 p.p.

2. Bramanti B, Thomas MG, Haak W, Unterlaender M, Jores P, Tambets K, et al. Genetic discontinuity between local hunter-gatherers and central Europe’s first farmers. Science. 2009;326(5949):137–40.

3. Brandt G, Haak W, Adler CJ, Roth C, Szecsenyi-Nagy A, Karimnia S, et al. Ancient DNA reveals key stages in the formation of central European mitochondrial genetic diversity. Science. 2013;342(6155):257–61.

4. Haak W, Balanovsky O, Sanchez JJ, Koshel S, Zaporozhchenko V, Adler CJ, et al. Ancient DNA from European early neolithic farmers reveals their near eastern affinities. PLoS biology. 2010;8(11):e1000536.

5. Haak W, Forster P, Bramanti B, Matsumura S, Brandt G, Tanzer M, et al. Ancient DNA from the first European farmers in 7500-year-old Neolithic sites. Science. 2005;310(5750):1016–8.

6. Gamba C, Fernandez E, Tirado M, Deguilloux MF, Pemonge MH, Utrilla P, et al. Ancient DNA from an Early Neolithic Iberian population supports a pioneer colonization by first farmers. Mol Ecol. 2012;21(1):45–56.

7. Allentoft ME, Sikora M, Sjogren KG, Rasmussen S, Rasmussen M, Stenderup J, et al. Population genomics of Bronze Age Eurasia. Nature. 2015;522(75):167–72.

8. Rasmussen M, Sikora M, Albrechtsen A, Korneliussen TS, Moreno-Mayar JV, Poznik GD, et al. The ancestry and affiliations of Kennewick Man. Nature. 2015;523(7561):455–8.

9. Olalde I, Schroeder H, Sandoval-Velasco M, Vinner L, Lobon I, Ramirez O, et al. A Common Genetic Origin for Early Farmers from Mediterranean Cardial and Central European LBK Cultures. Molecular biology and evolution. 2015.

10. Lazaridis I, Patterson N, Mittnik A, Renaud G, Mallick S, Kirsanow K, et al. Ancient human genomes suggest three ancestral populations for present-day Europeans. Nature. 2014;513(7518):409–13.

11. Gamba C, Jones ER, Teasdale MD, McLaughlin RL, Gonzalez-Fortes G, Mattiangeli V, et al. Genome flux and stasis in a five millennium transect of European prehistory. Nat Commun. 2014;5:5257.

12. Hervella M, Izagirre N, Alonso S, Fregel R, Alonso A, Cabrera VM, et al. Ancient DNA from hunter-gatherer and farmer groups from Northern Spain supports a random dispersion model for the Neolithic expansion into Europe. PloS one. 2012;7(4):e34417.

13. Bollongino R, Nehlich O, Richards MP, Orschiedt J, Thomas MG, Sell 2000 years of parallel societies in Stone Age Central Europe. Science. 2013;342(6157):479–81.

14. Deguilloux MF, Mendisco F. Ancient DNA: A window to the past of Europe. Human heredity. 2013;76(3-4):121–32.

15. Anderson CN, Ramakrishnan U, Chan YL, Hadly EA. Serial SimCoal: a population genetics model for data from multiple populations and points in time. Bioinformatics. 2005;21(8):1733–4.

16. Excoffier L, Foll M. fastsimcoal: a continuous-time coalescent simulator of genomic diversity under arbitrarily complex evolutionary scenarios. Bioinformatics. 2011;27(9):1332–4.

17. Hofmanová Z, Kreutzer S, Hellenthal G, Sell C, Diekmann Y, Díez del Molino D, et al. Early farmers from across Europe directly descended from Neolithic Aegeans. bioRxiv. 2015.

18. Manni F, Toupance B, Sabbagh A, Heyer E. New method for surname studies of ancient patrilineal population structures, and possible application to improvement of Y-chromosome sampling. American journal of physical anthropology. 2005;126(2):214–28.

19. Manni F, Barrai I. Genetic structures and linguistic boundaries in Italy: a microregional approach. Human biology. 2001;73(3):335–47.

20. Heyer E. Population structure and immigration; a study of the Valserine valley (French Jura) from the 17th century until the present. Annals of human biology. 1993;20(6):565–73.

21. Bideau A, Brunet G, Heyer E, Plauchu H, Robert JM. An abnormal concentration of cases of Rendu-Osler disease in the Valserine valley of the French Jura: a genealogical and demographic study. Annals of human biology. 1992;19(3):233–47.

22. Sanchez-Quinto F, Schroeder H, Ramirez O, Avila-Arcos MC, Pybus M, Olalde I, et al. Genomic affinities of two 7,000-year-old Iberian hunter-gatherers. Current biology: CB. 2012;22(16):1494–9.

23. Rasteiro R, Chikhi L. Female and male perspectives on the neolithic transition in Europe: clues from ancient and modern genetic data. PloS one. 2013;8(4):e60944.

24. Arenas M, Francois O, Currat M, Ray N, Excoffier L. Influence of admixture and paleolithic range contractions on current European diversity gradients. Molecular biology and evolution. 2013;30(1):57–61.

25. Currat M, Ruedi M, Petit RJ, Excoffier L. The hidden side of invasions: massive introgression by local genes. Evolution; international journal of organic evolution. 2008;62(8):1908–20.

26. Francois O, Currat M, Ray N, Han E, Excoffier L, Novembre J. Principal component analysis under population genetic models of range expansion and admixture. Molecular biology and evolution. 2010;27(6):1257–68.

27. Ray N, Currat M, Excoffier L. Intra-deme molecular diversity in spatially expanding populations. Molecular biology and evolution. 2003;20(1):76–86.

28. Ray N, Currat M, Foll M, Excoffier L. SPLATCHE2: a spatially explicit simulation framework for complex demography, genetic admixture and recombination. Bioinformatics. 2010;26(23):2993–4.

29. Currat M, Excoffier L. The effect of the Neolithic expansion on European molecular diversity. Proceedings Biological sciences / The Royal Society. 2005;272(1564):679–88.

30. Currat M, Ray N, Excoffier L. SPLATCHE: a program to simulate genetic diversity taking into account environmental heterogeneity. Mol Ecol Notes. 2004;4(1):139–42.

31. Zimmermann A, Hilpert J, Wendt KP. Estimations of population density for selected periods between the Neolithic and AD 1800. Human biology. 2009;81(2-3):357–80.

32. Excoffier L, Lischer HE. Arlequin suite ver 3.5: a new series of programs to perform population genetics analyses under Linux and Windows. Molecular ecology resources. 2010;10(3):564–7.

33. Alroy J. A multispecies overkill simulation of the end-Pleistocene megafaunal mass extinction. Science. 2001;292(5523):1893–6.

34. Steele J, Adams J, Sluckin T. Modelling Paleoindian dispersals (Paleoecology and human populations). World Archaeol. 1998;30(2):286–305.

35. Pinhasi R, Fort J, Ammerman AJ. Tracing the origin and spread of agriculture in Europe. PLoS biology. 2005;3(12):e410.

36. Wegmann D, Leuenberger C, Neuenschwander S, Excoffier L. ABCtoolbox: a versatile toolkit for approximate Bayesian computations. BMC Bioinformatics. 2010;11:116.

37. Slatkin M. Inbreeding coefficients and coalescence times. Genetical research. 1991;58(2):167–75.

38. Currat M, Excoffier L, Maddison W, Otto SP, Ray N, Whitlock MC, et al. Comment on “Ongoing adaptive evolution of ASPM, a brain size determinant in Homo sapiens"” and “Microcephalin, a gene regulating brain size, continues to evolve adaptively in humans”. Science. 2006;313(5784):172; author reply

39. Deshpande O, Batzoglou S, Feldman MW, Cavalli-Sforza LL. A serial founder effect model for human settlement out of Africa. Proceedings Biological sciences/TheRoyal Society. 2009;276(1655):291–300.

40. Slatkin M, Excoffier L. Serial Founder Effects During Range Expansion: A Spatial Analog of Genetic Drift. Genetics. 2012;191(1):171–81.

41. Wakeley J. Nonequilibrium migration in human history. Genetics. 1999;153(4):1863–71.

42. Wakeley J. The coalescent in an island model of population subdivision with variation among demes. Theor Popul Biol. 2001;59(2):133–44.

43. Bocquet-Appel J-P. The neolithic demographic transition and its consequences. Dordrecht: Springer; 2008. 542 p. p.

44. Excoffier L. Patterns of DNA sequence diversity and genetic structure after a range expansion: lessons from the infinite-island model. Molecular ecology. 2004;13(4):853–64.

45. Mathieson I, Lazaridis I, Rohland N, Mallick S, Patterson N, Roodenberg SA, et al. Genome-wide patterns of selection in 230 ancient Eurasians. Nature. 2015;528(7583):499–503.

46. Omrak A, Gunther T, Valdiosera C, Svensson EM, Malmstrom H, Kiesewetter H, et al. Genomic Evidence Establishes Anatolia as the Source of the European Neolithic Gene Pool. Current biology: CB. 2016;26(2):270–5.

47. Haak W, Lazaridis I, Patterson N, Rohland N, Mallick S, Llamas B, et al. Massive migration from the steppe was a source for Indo-European languages in Europe. Nature. 2015;522(7555):207.

48. Brotherton P, Haak W, Templeton J, Brandt G, Soubrier J, Adler CJ, Neolithic mitochondrial haplogroup H genomes and the genetic origins of Europeans. Nature communications. 2013;4:1764.

49. Fu Q, Mittnik A, Johnson PLF, Bos K, Lari M, Bollongino R, et al. A revised timescale for human evolution based on ancient mitochondrial genomes. Current Biology. 2013;23(7):553–9.

